# Generating realistic cell samples for gene selection in scRNA-seq data: A novel generative framework

**DOI:** 10.1101/2021.04.29.441920

**Authors:** Snehalika Lall, Sumanta Ray, Sanghamitra Bandyopadhyay

## Abstract

High dimensional, small sample size (HDSS) scRNA-seq data presents a challenge to the gene selection task in single cell. Conventional gene selection techniques are unstable and less reliable due to the fewer number of available samples which affects cell clustering and annotation. Here, we present an improved version of generative adversarial network (GAN) called LSH-GAN to address this issue by producing new realistic samples and combining this with the original scRNA-seq data. We update the training procedure of the generator of GAN using locality sensitive hashing which speeds up the sample generation, thus maintains the feasibility of applying gene selection procedures in high dimension scRNA-seq data. Experimental results show a significant improvement in the performance of benchmark feature (gene) selection techniques on generated samples of one synthetic and four HDSS scRNA-seq data. Comprehensive simulation study ensures the applicability of the model in the feature (gene) selection domain of HDSS scRNA-seq data.

**Availability:** The corresponding software is available at https://github.com/Snehalikalall/LSH-GAN

## 1 Introduction

Identifying essential features is a persistent problem in machine learning, which is generally known as the feature selection problem [1]. Recently, the emergence of high dimensional biological data such as single cell RNA sequence (scRNA-seq) data has posed a significant challenge to the machine learning researchers [2, 3]. Handling the high dimension, and small sample size (HDSS) data is difficult for feature selection (FS) which posed a problem in classification techniques. Particularly, it affects both the accuracy of the classification and increases the risk of overfitting. A few outliers can drastically affect the FS techniques, and the selected feature sets may not be adequate to discriminate the classes [4]. Moreover, high dimensionality increases the computational time beyond acceptablity. So, feature selection is essential to reduce the dimensionality of the data for further processing.

HDSS data is prevalent in the single cell domain due to the budgetary constraint of single cell experiments. The general pipeline of scRNA-seq downstream analysis starts with pre-processing (normalization, quality control) of the raw count matrix and then going through several steps which include identification of relevant genes, clustering of cells, and annotating cell clusters with marker genes [5–9]. Each step has a profound effect on the next stage of analysis. The gene selection step identifies the most relevant features/genes from the normalized/preprocessed data and has an immense impact on the cell clustering. The general procedure for selecting relevant genes which are primarily based on high variation (highly variable genes) [10, 11] or significantly high expression (highly expressed genes) [5] suffers from a small sample effect. The general FS techniques also failed to provide a stable and predictive feature set in this case due to an ultra large size of feature (gene). One way to solve this issue is to go for a robust and stable technique that does not overfit the data. A few attempts [12–14] were observed recently which embed statistical and information-theoretic approach. Although these methods result in stable features, however, these are not performed well in small sample scRNA-seq data.

In this paper, we propose a generative model to sort out the problem of feature (gene) selection in HDSS scRNA-seq data. It can be noted that if the sample size is sufficiently large, the selected feature sets have a high probability of containing the most relevant and discriminating features. We use generative adversarial model to generate more samples to balance between feature and sample size. Generative Adversarial Network (GAN) [15] has already been shown to be a powerful technique for learning and generating complex distributions [16, 17]. However, the training procedure of GAN is difficult and unstable. The training suffers from instability because both the generator and the discriminator model are trained simultaneously in a game that requires a Nash equilibrium to complete the procedure. Gradient descent does this, but sometimes it doesn’t, which results in a costly time consuming training procedure. The main contribution here is in modifying the generator input that results in a fast training procedure. We create a subsample of original data based on locality sensitive hashing (LSH) technique and augment this with noise distribution, which is given as input to the generator. Thus, the generator does not take pure noise as input, instead, we introduce a bias in it by augmenting a subsample of data with the noise distribution.

Researchers are still trying to find improved versions of the generative adversarial network (GAN) to use in different domains. Most of the variations such as progressive GAN (PGAN) [18], Wasserstein GAN (WGAN) [16] try to train the model quicker than the conventional GAN. Unlike PGAN and WGAN, conditional GAN (CGAN) [19] operates by conditioning the conventional model on additional data sources (maybe class label or data from different modalities) to dictate the data generation. In our model, we direct our attention to the additional sample generation from HDSS data. However, the generated sample size becomes increasingly large with more features, the generation of which may not be feasible for conventional generative models. Augmenting subsample of real data distribution (*p_data_*(*x*)) with the prior noise (*p_z_*(*z*)) makes the training procedure of our model faster than the conventional GAN. We theoretically proved that the global minimum value of the virtual training criterion of the generator is less than the traditional GAN (< -*log4*).

### Summary of contributions

Here, we provide the following novelties:

– The proposed model is the first one to address the problem of gene selection in HDSS scRNA-seq data using generative model.
– LSH-GAN can able to generate realistic samples in a faster way than the traditional GAN. This makes LSH-GAN more feasible to use in the feature (gene) selection problem of scRNA-seq data.
– Here we derive a new way of training a generator that combines subsamples of original data with pure noise and takes this as input.
– For a fixed number of iteration LSH-GAN performed better than the traditional GAN in generating realistic samples.
– Gene selection on the generated samples of LSH-GAN provides excellent results for small-sample and large-feature sized single cell data.

## 2 Method

In this section, we first provide a short description of Generative Adversarial Network (GAN) and then describe our proposed LSH-GAN model. Later, we explain the theoretical foundation of LSH-GAN model, and finally provide its application in the gene selection task of scRNA-seq HDSS data.

### 2.1 Generative adversarial network

Generative adversarial network (GAN) is introduced in [15] which was proposed to train a generative model. GAN consists of two blocks: a generative model (*G*) that learn the data distribution (*p*(*x*)), and a discriminative model (*D*) that estimates the probability that a sample came from the training data (*X*) rather than from the generator (*G*). These two models can be non-linear mapping functions such as two neural networks. To learn the generator distribution *p_g_* over data *x*, a differentiable mapping function is built by generator *G* to map a prior noise distribution *p_z_*(*z*) to the data space as *G*(*z*; *θ_g_*). The discriminator function *D*(*x*; *θ_d_*) returns a single scalar that represents the probability of *x* coming from the real data rather than from generator distribution *p_g_*. The goal of the generator is to fool the discriminator, which tries to distinguish between true and generated data. Training of *D* ensures that the discriminator can properly distinguish samples coming from both training samples and the generator. *G* and *D* are simultaneously trained to minimize *log*(1 — *D*(*G*(*z*)) for *G* and maximize *log*(*D*(*x*)) for *D*. It forms a two-player min-max game with value function *V*(*G*,*D*)

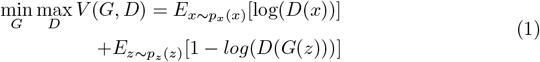

In the following, we will describe the workflow of our analysis pipeline.

### 2.2 Proposed model: LSH-GAN

The figure 1 describes the workflow of our analysis pipeline. Figure 1, panel-A, describes the application of the proposed LSH-GAN model in feature selection task of the HDSS scRNA-seq data data, while Panel-B depicts basic building blocks of the model. The following subsections describe in brief.

**Fig 1.**
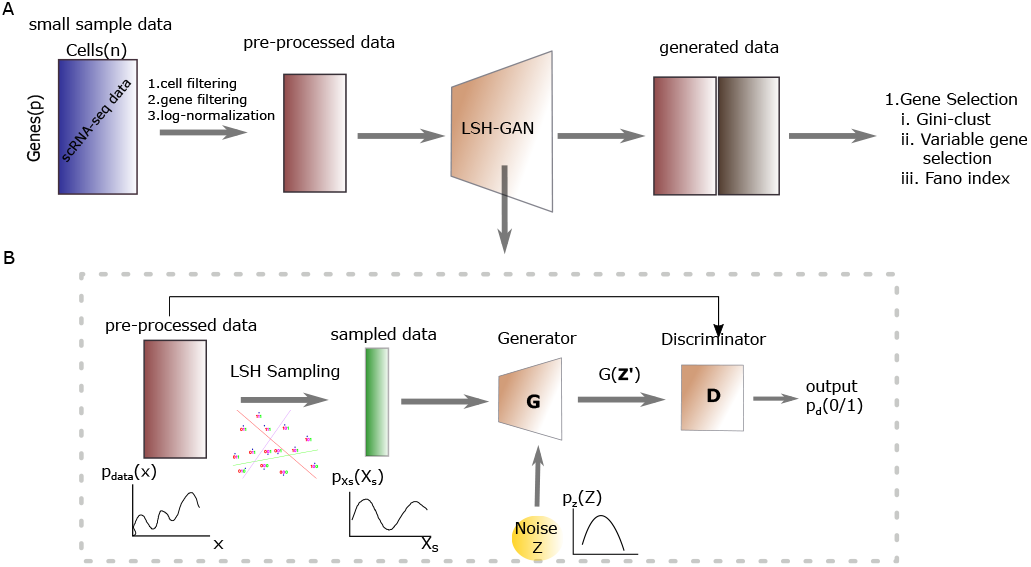
Panel-A: Figure shows the workflow for gene selection in HDSS scRNA-seq data using generated samples with LSH-GAN model. Panel-B shows the general architecture of LSH-GAN.

#### LSH step: sampling of input data

Locality Sensitive Hashing (LSH) [13, 20, 21] is widely used in nearest neighbor searching to reduce the dimensionality of data. LSH utilizes locality-sensitive hash functions which hash similar objects into the same bucket with a high probability. The number of buckets is much lesser than the universe of possible items, thus reduces the search space of the query objects (see supplement for detailed description of LSH technique).

In this work first, the unique hash codes which depict the local regions or neighborhood of each data point are produced. For this, we utilized python sklearn implementation of *LSHForest* module with default parameters.An approximate neighborhood graph (*κ*-nn graph) is constructed by using *κ* = 5 for each data point. This step computes the euclidean distances between the query point and its candidate neighbors. Sampling is carried out in a ‘greedy’ fashion where each data point is *traversed* sequentially and its corresponding five nearest neighbors are flagged out which never visited again. Thus after one traversing a sub-set of samples is obtained which is further down-sampled by performing the same step iteratively.

#### Generator of LSH-GAN

The generator function (*G*) is modified by augmenting its taken input data. Instead of giving the pure noise (*p_z_*(*z*)) as input we augment a subsample of real data distribution (*p_data_*(*x*)) with it. The sampling of the input data is done in the LSH step. Thus the Generator (G) function builds a mapping function from 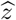 to data space (*x*) as 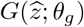 and is defined as: 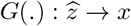. Modifying the generator in this way we claim that it can increase the probability of generating samples of real data in lesser time.

#### Discriminator of LSH-GAN

Here discriminator (D) takes both the real data *p_data_*(*x*) and generated data coming from generator 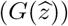, with probability density 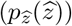 and returns the scalar value, *D*(*x*) that represents the probability that the data *x* is coming form the real data: *D*(.): *x* → [0,1].

So, the value function can be written as:

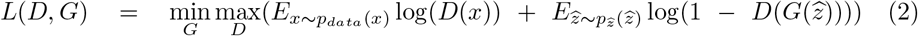

D and G forms a two-player minimax game with value function *L*(*G*, *D*). We train *D* to maximize the probability of correctly validate the real data and generated data. We simultaneously train *G* to minimize 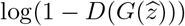, where 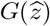 represents the generated data from the generator by taking the noise (*p_z_*) and the sampled data *p_X_s__* (*x_s_*) as input.

#### Feature/gene selection using LSH-GAN

The generated cell samples are utilized for gene selection task. We have employed three feature selection methods (*CV*^2^ *Index*, *pca-loading* and *Fano Factor*), widely used for the gene selection task in scRNA-seq data. Single cell clustering method (SC3) technique is utilized to validate the selected genes from the generated samples.

The whole algorithm and the sampling procedure is described in the ‘LSH-GAN algorithm’.

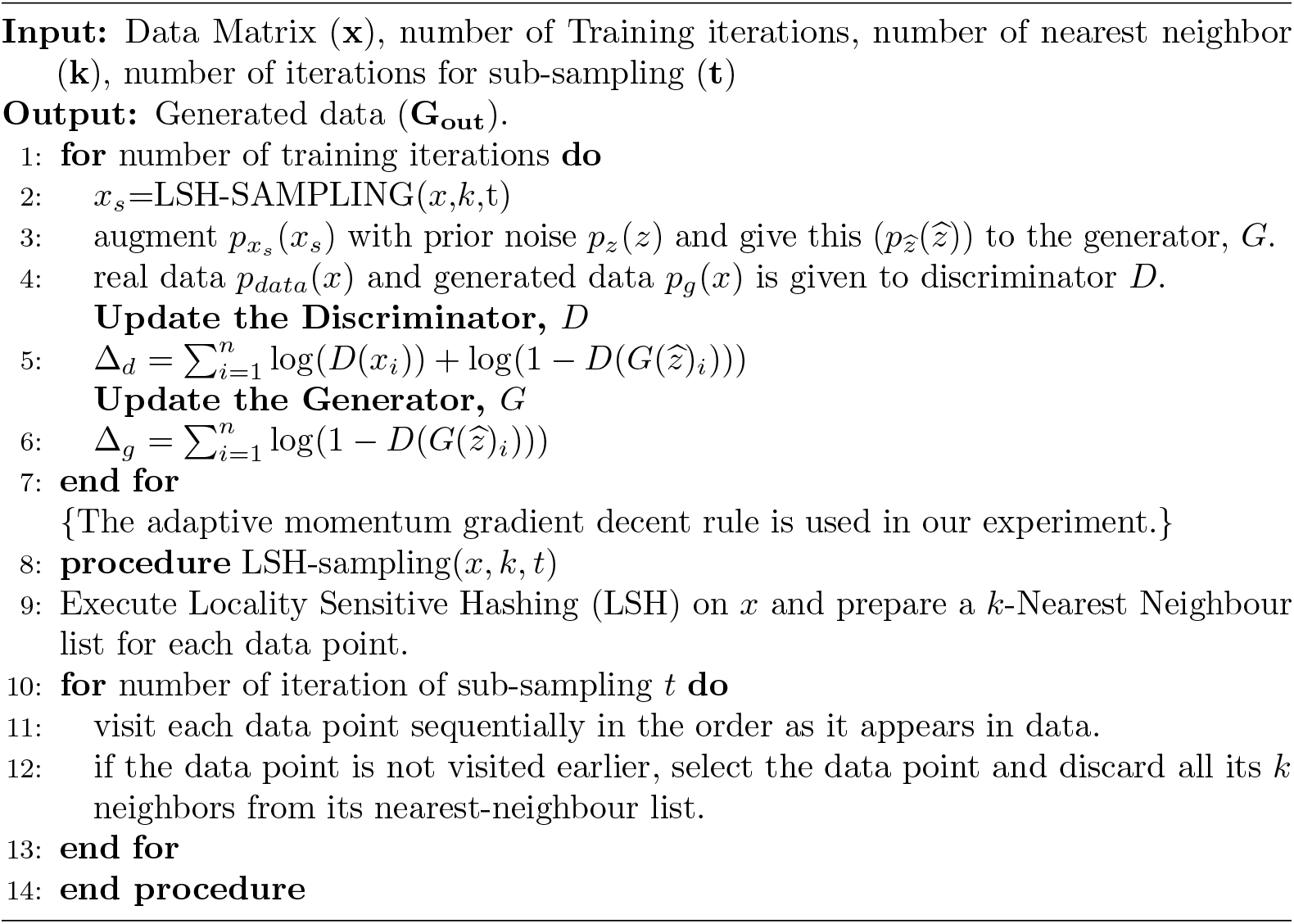

### 2.3 Theoretical Analysis of LSH-GAN

Following the above section, a sub-sampling of real data *p_X_s__* (*x_s_*) is augmented with the prior noise distribution, *p_z_*(*z*). Due to this additional information in generator, we assume that the probability 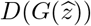 will increase by a factor, ζ.

**Proposition**. *L*(*D*, *G*) is maximized with respect to Discriminator (*D*), for a fixed Generator (*G*), when

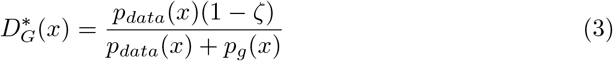

**Proof**. Equation 2 can be written as:

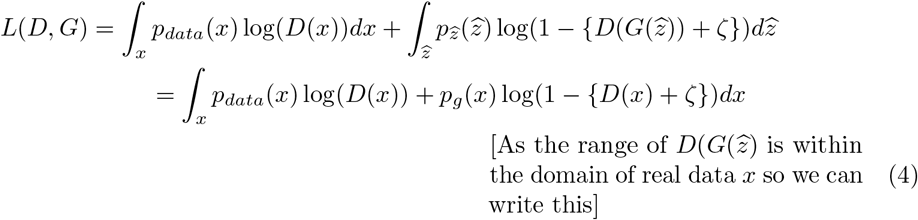

We know that, the function *y* = *a* log *x* + *b* log(1 — (*x* + ζ)) will have maximum value, at 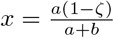, for any (*a*, *b*) ∈ *R*^2^{0,0} and ζ ∈ (0,1). So, the optimum value of *D* for a fixed generator, *G* is:

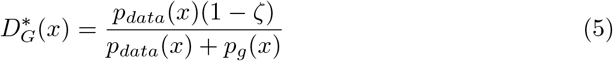

The training objective for discriminator *D* is to maximize the log-likelihood of the conditional probability *P*(*Y* = *y*|*x*), where *Y* signify whether *x* is coming from real data distribution(*y* = 1) or coming from the generator(*y* = 0). Now the equation 2 can be written as:

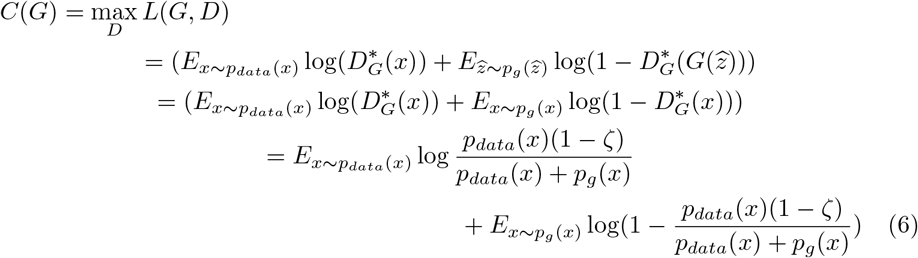

**Theorem** At *p_g_*(*x*) = *p_data_*(*x*) (global minimum criterion of value function *L*(*G*, *D*)), the value of *C*(*G*) is less than (—log 4).

**proof** From equation 6 we get

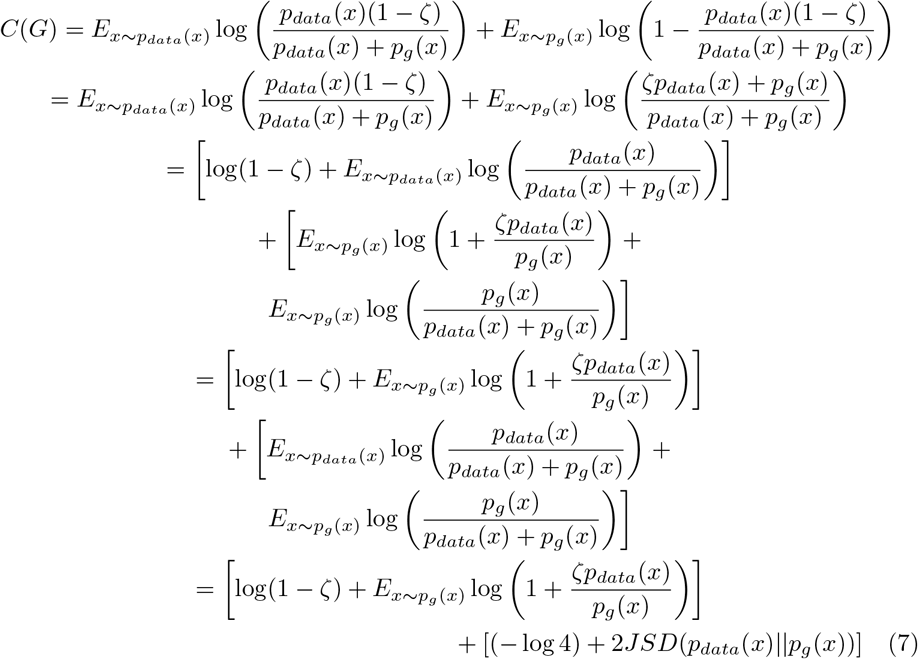

where, *JSD*(*p_data_*(*x*)||*p_g_*(*x*)) represents Jensen–Shannon divergence between two distributions *p_data_* and *p_g_*. Now, if the two distribution are equal, Jensen–Shannon divergence (JSD) will be zero. Thus, for global minimum criterion of the value function (*p_g_* = *p_data_*) the Equation 7 is reduces to,

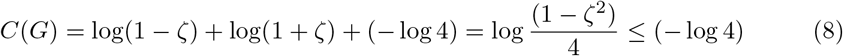

This completes the proof.

## 3 Results

For experimental validation first, we validate our proposed model in synthetic data. The aim is to see the performance of LSH-GAN in generating realistic samples. Next, we validate whether the generated samples can be used in the feature selection task. Finally, we apply the proposed model on HDSS scRNA-seq data and use benchmark gene selection methods on the generated samples. In both datasets, we compare the performance of LSH-GAN with the traditional GAN.

### 3.1 Dataset

We have used four public benchmark scRNA-seq datasets: Pollen [22], Darmanis [23], Yan [24] and Klein [25], downloaded from Gene Expression Omnibus (GEO) https://www.ncbi.nlm.nih.gov/geo/. Table 1 shows a detailed summary of the used datasets (see supplement for description). The sample:feature ratio for all the datasets are less than 0.012.

**Table 1.**
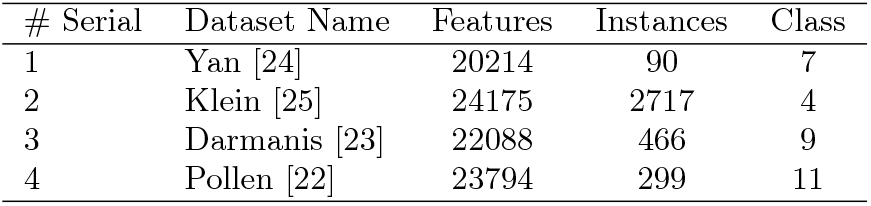
A brief summary of the datasets used in the experiments.

### 3.2 Experimental settings

The number of nearest neighbor (*κ*) and the number of iteration (*t*) are two main parameters of the LSH-step (see Algorithm), tuning of which affects the amount of sampling given to the generator for training the LSH-GAN model. We vary *κ* and *t* in the range {5,10,15, 20} and {1, 2}, respectively, and choose that value for which the Wasserstein distance [16] between generated and real samples is reported to be minimum. We fixed the amount of sampling using *κ* = 5, *t* = 1 for Pollen, Yan, Darmanis datasets and *κ* = 5, *t* = 2 for Klein dataset. For generating hash code from LSH sampling, *LSHForest* of *scikit-learn* version 0.19.2 is utilized.

Tuning of another parameter, *S_gen_* which represents the size of generated samples is required for the feature selection step. Here *S_gen_* is set within the range {0.25p, 0.5p, 0.75p, 1p, 1.25p, 1.5p}, where *p* is the feature number. We take the adaptive learning rate optimization algorithm implemented in ADAM optimizer in python Tensorflow version 1.9.2. Generator (G) and Discriminator (D) uses 2-layer multilayer perceptrons with hidden layer dimension as (16, 16). For traditional GAN, we retain the same settings as LSH-GAN for G and D networks.

Three well known gene selection methods (with default parameters) of scRNA-seq data are adopted for validation: *PCA_Loading* [26], *CV*^2^ *Index* [27] and *Fano Factor* [28]. We select top 100 features (genes) using all three feature selection methods on synthetic and scRNA-seq datasets. For validation purposes, Wasserstein metric [16] is utilized to estimate the quality of the generated data. Clustering of scRNA-seq data is performed using *SC*3 [29] technique with default parameters. Clustering performance is evaluated using the Adjusted Rand Index (ARI).

### 3.3 Data Preprocessing

The raw count matrix 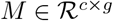, where *c* and *g* represents the number of cells and genes, respectively, is normalized using *Linnorm* [30] Bioconductor package of R. We select cells having more than a thousand expressed genes (non zero values) and choose genes having a minimum read count more than 5 in at least 10% of the cells. *log_2_* normalization is performed on the transformed matrix by adding one as a pseudo count.

### 3.4 LSH-GAN generates samples faster than traditional GAN on HDSS synthetic data

We train the LSH-GAN on HDSS synthetic data and generate realistic samples to compare against the traditional GAN model. For this, we create a 2-class non-overlapping Gaussian mixture data consisting 100 samples and 1000 features by taking the mean (*μ*) of the data in the range of 5 to 15 for class-1 and —15 to —5 for class-2. The covariance matrix (*Σ*) is taken for all the samples using the formula 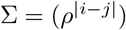, where *i*, *j* are row and column index, and *ρ* is equal to 0.5. We calculate Wasserstein distance between the real data distribution (*p_data_*) and the generated data distribution (*p_g_*) to estimate the quality of the generated data.

We use different settings of *k^th^* (*κ*=5, 10, 15, 20) nearest neighbor to generate sub-sample of data from LSH sampling procedure. In each case, the sampled data (*p_X_s__*) is augmented with prior noise (*p_z_*) and given to the generator of LSH-GAN for model training.

For comparison with the traditional GAN model, we use the data with train: test split of 80:20 and calculate the Wasserstein metric between the test sample and the generated sample. Table 2 shows the values of the metric for LSH-GAN and traditional GAN model in different range of epochs and nearest neighbors *κ*. A closer look into the table 2 reveals that the performance of LSH-GAN (at 10000 epoch and *κ* = 5) is far better than the traditional GAN model with 25000 epochs. Notably, for less amount of sampling (larger *κ*), LSH-GAN needs more iterations for training. As for particular example, the performance of LSH-GAN achieved on 20000 epoch and *κ* = 20 is rivaled only at 10000 epoch for *κ* = 10. Thus it is evident from the results that reducing the amount of sampling needs more epochs and thus needs more training time for the LSH-GAN model to converge. Figure 2 also supports this statement. Here, the two models (LSH-GAN, and traditional GAN) are trained to simulate a two dimensional synthetic data of known distribution, for which the LSH-GAN can able to generate samples that are more real than the traditional GAN, in a lesser number of iteration.

**Fig 2.**
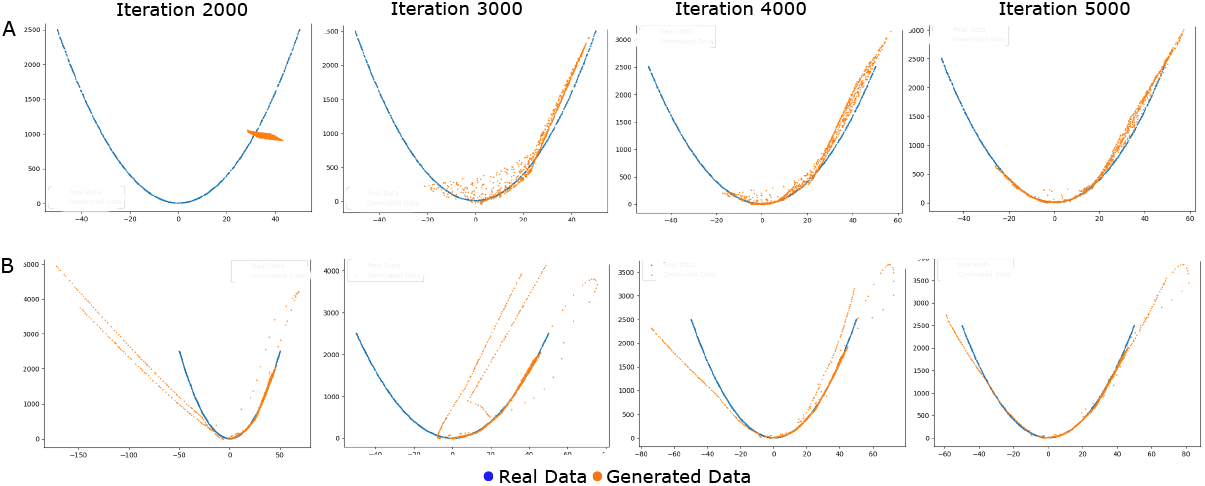
Generation of two dimensional synthetic data using traditional GAN (upper row, Panel-A) and LSH-GAN (lower row, Panel-B) model for different epochs.

**Table 2.** Wasserstein distance between generated and real data distribution. Model is trained on synthetic data of size 100 × 1000 Gaussian mixture data with 2 non-overlapping classes.

| Nearest Neighbour | Model | Epoch |  |  |  |
| --- | --- | --- | --- | --- | --- |
|  |  | 10000 | 15000 | 20000 | 25000 |
| $k = 5$ | LSH-GAN | 0.46 | 0.35 | 0.33 | 0.45 |
| $k = 10$ | LSH-GAN | 1.09 | 0.89 | 0.83 | 0.82 |
| $k = 15$ | LSH-GAN | 1.36 | 0.89 | 1.45 | 0.87 |
| $k = 20$ | LSH-GAN | 1.53 | 1.35 | 1.19 | 0.83 |
|  | GAN | 1.71 | 1.73 | 1.75 | 1.70 |

### 3.5 Gene selection in HDSS scRNA-seq data using LSH-GAN

We trained the LSH-GAN model in four small sample scRNA-seq data (see table 1). Here, a sub-sample of real data distribution is augmented with prior noise and used as the input to the generator network. The generated data using LSH-GAN (with *κ*=5) is validated by computing the Wasserstein metric between the real and generated data distribution for different epochs (see figure 3). For each data, we note the epoch (*e_opt_*), which results in the lowest Wasserstein metric. For example, we take *e_opt_* as 10k, 30k, 10k, and 15k for the dataset Darmanis, Yan, Pollen, and Klein, respectively.

**Fig 3.**
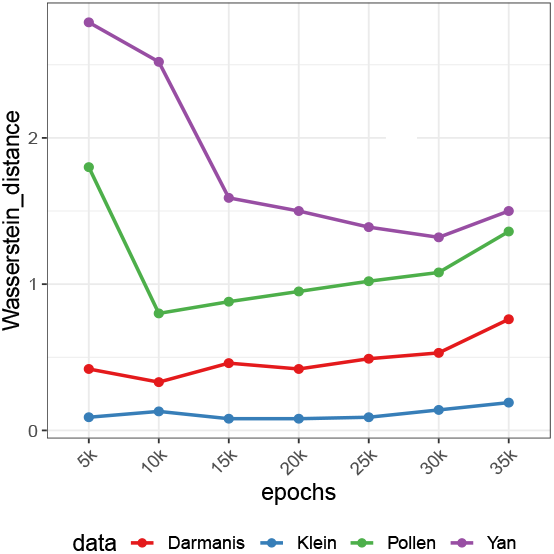
Figure shows Wasserstein metric between real and generated data distribution across different epochs for scRNA-seq datasets.

Here, we have employed three feature selection methods (*CV*^2^ *Index*, *pca-loading* and *Fano Factor*), widely used for the gene selection task in scRNA-seq data and one single cell clustering method (SC3) technique to validate the selected genes from the generated data. To know the correct amount sampling, we select the optimal sample size *S_opt_* by observing the ARI values of cell clustering results for different sample sizes *S_gen_*={0.25p, 0.5p, 0.75p, 1p, 1.5p}. Figure 4 shows the results of clustering using the features selected with the three gene selection techniques on four scRNA-seq datasets. A closer look in the figure 4 reveals that, for most of the gene selection techniques, sample size (*S_opt_*) 1.25*p* and 1.5*p* (*p* denotes number of features) results best ARI values.

**Fig 4.**
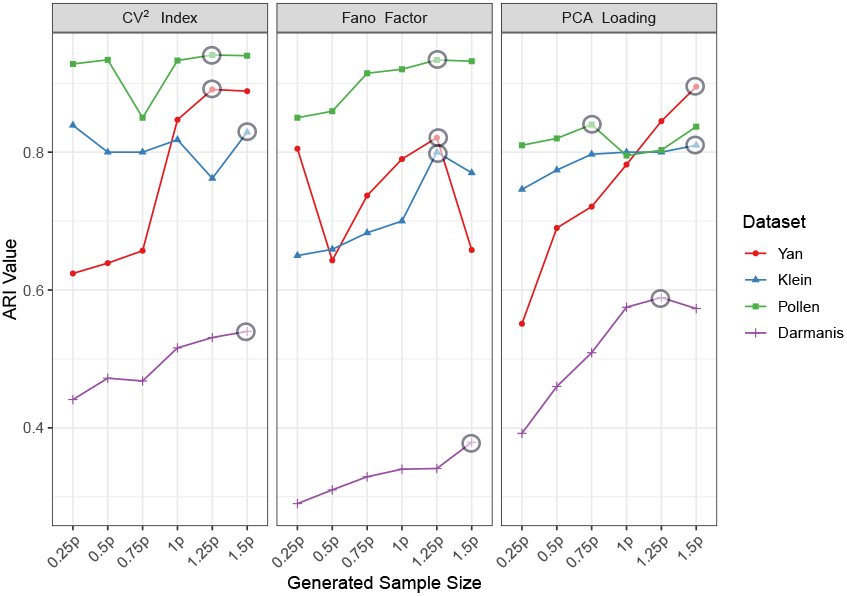
Figure shows the line plots signifying the variation of ARI values of clustering results with different sizes of generated samples. SC3 clustering is performed using the selected genes with three different gene selection techniques. Gene selection is performed on the generated data with sample:feature ratio as 0.25, 0.5, 0.75, 1, 1,25 and 1.5. Higher ARI values are marked as encircled points in each line plot.

LSH-GAN is compared with traditional GAN in the gene selection problem of HDSS scRNA-seq datasets. LSH-GAN is trained with the selected epoch (*e_opt_*) and optimal sample size (*S_opt_*) for generating samples from the scRNA-seq data. The aim is to know whether the selected features/genes from the generated combined data can lead to a pure clustering of cells. Table 3 shows the comparisons of the ARI values resulting from the cell clustering. It is evident from the table that features/genes selected from the generated combined data of the LSH-GAN model (with *e_opt_* and *S_opt_*) produces better clustering results than the traditional GAN model. Here, two models are trained with the same number of epochs.

**Table 3.** Table shows the comparison (ARI values) between GAN and LSH-GAN for feature selection in HDSS scRNA-seq data.

| Data | epoch<br>($e_{opt}$) | FS method | Sample size($S_{opt}$) | LSH-GAN | GAN | without<br>model |
| --- | --- | --- | --- | --- | --- | --- |
| Darmanis | 10k | PCA_Loading | 1.25p | 0.589 | 0.129 | 0.4 |
|  |  | Fano Factor | 1.5p | 0.435 | 0.27 | 0.34 |
| | | $CV^2$ index | 1.5p | 0.598 | 0.461 | 0.457 |
| Yan | 30k | PCA_Loading | 1.5p | 0.895 | 0.62 | 0.66 |
|  |  | Fano Factor | 1.25p | 0.821 | 0.8 | 0.793 |
| | | $CV^2$ index | 1.25p | 0.891 | 0.719 | 0.80 |
| Pollen | 10k | PCA_Loading | 1.5p | 0.835 | 0.788 | 0.80 |
|  |  | Fano Factor | 1.25p | 0.933 | 0.915 | 0.712 |
| | | $CV^2$ index | 1.25p | 0.94 | 0.906 | 0.81 |
| Klein | 15k | PCA_Loading | 1.5p | 0.815 | 0.581 | 0.66 |
|  |  | Fano Factor | 1.25p | 0.8 | 0.466 | 0.796 |
| | | $CV^2$ index | 1.5p | 0.82 | 0.48 | 0.68 |

### 3.6 Selected genes using LSH-GAN can effectively predict cell clusters

Here we provide the detailed results of clustering on four used datasets using the genes selected from the generated samples. For this, we adopted a widely used single cell clustering method SC3 [29]. Figure 5 panel-A, depicts the t-SNE visualization of predicted clusters and their original labels for Yan and Pollen datasets (see supplementary figure-1 for the results of other two datasets). Panel-B of figure 5 represents heatmaps of cell × cell consensus matrix. Each heatmap signifies the number of times a pair of cells is appearing in the same cluster [29]. Here two cells are said to be in different clusters if the score is zero (blue color). Similarly, a score ‘1’ (red) signifies two cells are belonging to the same class. Thus a completely red diagonals and blue off-diagonals represent a perfect clustering. A careful notice on the figure 5, panel-A and -B reveals a perfect match between the original and predicted labels for YAN and Pollen datasets.

**Fig 5.**
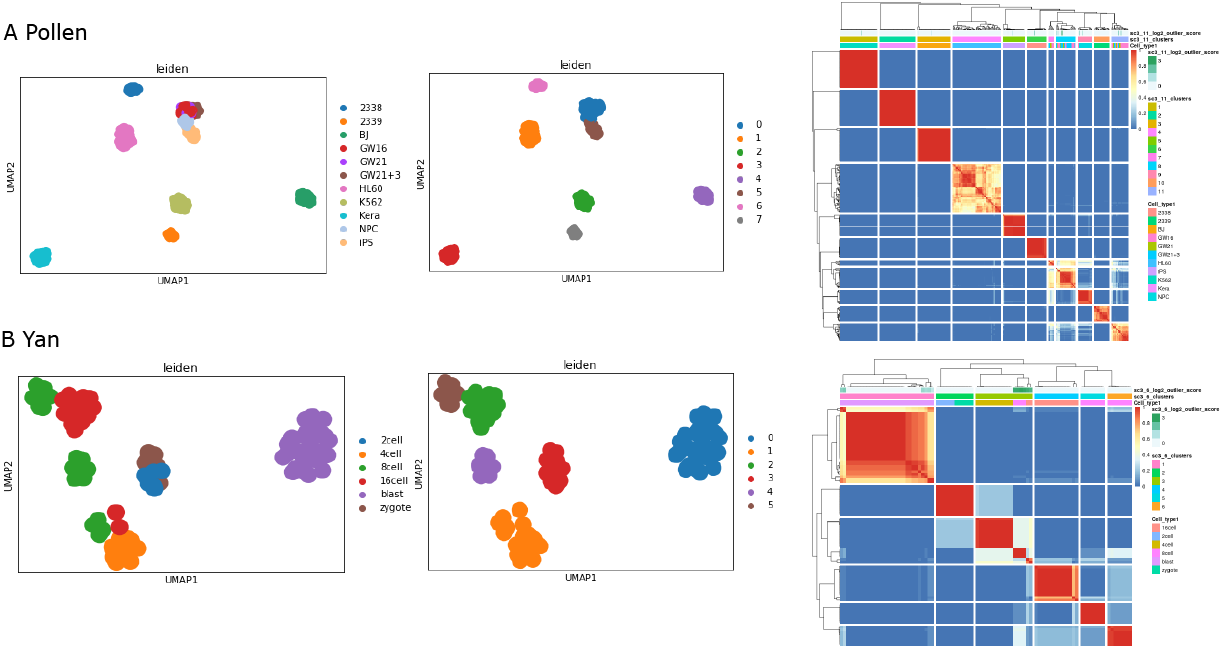
Figure shows the clustering results of Pollen and Yan data sets. Panel-A shows the t-SNE visualization of clustering results (original and predicted labels), whereas panel-B shows the consensus clustering plots of obtained clusters.

## 4 Discussions

In this paper, we present a novel and faster way of generating cell sample of HDSS single cell RNA-seq data using a generative model called LSH-GAN. We update the training procedure of generative adversarial network (GAN) using locality sensitive hashing which can produce realistic samples in a lesser number of iteration than the traditional GAN model. We utilized this in a valid problem of gene selection in HDSS single cell data. Our preliminary simulation experiment suggests that for a fixed number of training iteration the proposed model can generate more realistic samples than the traditional GAN model. This observation is also established theoretically by proving that the cost of value function is less than —*log*4 which is the cost for traditional GAN at the global minimum of virtual training criterion (*p_g_* = *p_data_*).

We demonstrated that the generated samples of LSH-GAN is useful for gene selection in HDSS scRNA-seq data. For validation of the generated cell samples, we use the conventional steps of downstream analysis for scRNA-seq data. We employ three widely used gene selection techniques and one single cell clustering technique for gene selection and grouping of cells using the selected genes. The precise clustering of cells demonstrates the quality of generated cell samples using the LSH-GAN model.

One limitation of our method is that for feature selection we hardly found any linear relationship between the clustering results with the sample size of generated scRNA-seq data. The correct sample size should be selected by using a different range of values between 0.25*p* to 1.5*p*, where *p* is the feature size. There may be some effects of different parameters related to single cell clustering (*SC*3 method) and feature selection (e.g. different FS method, number of selected features, etc.) which may play a critical role in the clustering performance. However, we found clustering results are always better for the generated data with more than 1p (p is the feature size) sample size. This observation suggests that for feature selection in HDSS data, whenever we produce samples larger than the feature size we will end up with a better clustering. The feasibility of generating such samples is justified by the faster training procedure of LSH-GAN model.

Taken together, the proposed model can generate good quality cell samples from HDSS scRNA-seq data in a lesser number of iteration than the traditional GAN model. Results show that LSH-GAN not only leads in the cell sample generation of scRNA-seq data but also accelerates the way of gene selection and cell clustering in the downstream analysis. We believe that LSH-GAN may be an important tool for computational biologists to explore the realistic cell samples of HDSS scRNA-seq data and its application in the downstream analysis.

## References

1. Cai J, Luo J, Wang S, Yang S. Feature selection in machine learning: A new perspective. Neurocomputing. 2018;300:70–79.

2. Vallejos CA, Risso D, Scialdone A, Dudoit S, Marioni JC. Normalizing single-cell RNA sequencing data: challenges and opportunities. Nature methods. 2017;14(6):565.

3. Ray S, Schonhuth A. MarkerCapsule: Explainable Single Cell Typing using Capsule Networks. bioRxiv. 2020;.

4. Liao S, Gao Q, Nie F, Liu Y, Zhang X. Worst-Case Discriminative Feature Selection. In: IJCAI; 2019. p. 2973–2979.

5. Duò A, Robinson MD, Soneson C. A systematic performance evaluation of clustering methods for single-cell RNA-seq data. F1000Research. 2018;7.

6. Luecken MD, Theis FJ. Current best practices in single-cell RNA-seq analysis: a tutorial. Molecular systems biology. 2019;15(6):e8746.

7. Ji Z, Ji H. TSCAN: Pseudo-time reconstruction and evaluation in single-cell RNA-seq analysis. Nucleic acids research. 2016;44(13):e117–e117.

8. Plass M, Solana J, Wolf FA, Ayoub S, Misios A, Glažar P, et al. Cell type atlas and lineage tree of a whole complex animal by single-cell transcriptomics. Science. 2018;360(6391).

9. Fincher CT, Wurtzel O, de Hoog T, Kravarik KM, Reddien PW. Cell type transcriptome atlas for the planarian Schmidtea mediterranea. Science. 2018;360(6391).

10. Chen G, Ning B, Shi T. Single-cell RNA-seq technologies and related computational data analysis. Frontiers in genetics. 2019;10:317.

11. Butler A, Hoffman P, Smibert P, Papalexi E, Satija R. Integrating single-cell transcriptomic data across different conditions, technologies, and species. Nature biotechnology. 2018;36(5):411–420.

12. Vans E, Patil A, Sharma A. FEATS: Feature selection based clustering of single-cell RNA-seq data. bioRxiv. 2020;.

13. Lall S, Sinha D, Bandyopadhyay S, Sengupta D. Structure-Aware Principal Component Analysis for Single-Cell RNA-seq Data. Journal of Computational Biology. 2018;.

14. Lall S, Ray S, Bandyopadhyay S. RgCop-A regularized copula based method for gene selection in single cell rna-seq data. bioRxiv. 2020;.

15. Goodfellow I, Pouget-Abadie J, Mirza M, Xu B, Warde-Farley D, Ozair S, et al. Generative adversarial nets. In: Advances in neural information processing systems; 2014. p. 2672–2680.

16. Arjovsky M, Chintala S, Bottou L. Wasserstein Generative Adversarial Networks. In: Proceedings of the 34th International Conference on Machine Learning. vol. 70 of Proceedings of Machine Learning Research. PMLR; 2017. p. 214–223.

17. Nowozin S, Cseke B, Tomioka R. f-gan: Training generative neural samplers using variational divergence minimization. arXiv preprint arXiv:160600709. 2016;.

18. Karras T, Aila T, Laine S, Lehtinen J. Progressive Growing of GANs for Improved Quality, Stability, and Variation. In: International Conference on Learning Representations; 2018. Available from: https://openreview.net/forum?id=Hk99zCeAb.

19. Mirza M, Osindero S. Conditional generative adversarial nets. arXiv preprint arXiv:14111784. 2014;.

20. Pauleve L, Jegou H, Amsaleg L. Locality sensitive hashing: A comparison of hash function types and querying mechanisms. Pattern Recognition Letters. 2010;31(11):1348–1358.

21. Mao XL, Feng BS, Hao YJ, Nie L, Huang H, Wen G. S2JSD-LSH: A locality-sensitive hashing schema for probability distributions. In: Proceedings of the AAAI Conference on Artificial Intelligence. vol. 31; 2017.

22. Pollen AA, Nowakowski TJ, Shuga J, Wang X, Leyrat AA, Lui JH, et al. Low-coverage single-cell mRNA sequencing reveals cellular heterogeneity and activated signaling pathways in developing cerebral cortex. Nature biotechnology. 2014;32(10):1053.

23. Darmanis S, Sloan SA, Zhang Y, Enge M, Caneda C, Shuer LM, et al. A survey of human brain transcriptome diversity at the single cell level. Proceedings of the National Academy of Sciences. 2015;112(23):7285–7290.

24. Yan L, Yang M, Guo H, Yang L, Wu J, Li R, et al. Single-cell RNA-Seq profiling of human preimplantation embryos and embryonic stem cells. Nature structural & molecular biology. 2013;20(9):1131.

25. Klein AM, Mazutis L, Akartuna I, Tallapragada N, Veres A, Li V, et al. Droplet barcoding for single-cell transcriptomics applied to embryonic stem cells. Cell. 2015;161(5):1187–1201.

26. Macosko EZ, Basu A, Satija R, Nemesh J, Shekhar K, Goldman M, et al. Highly parallel genome-wide expression profiling of individual cells using nanoliter droplets. Cell. 2015;161(5):1202–1214.

27. Brennecke P, Anders S, Kim JK, Kołodziejczyk AA, Zhang X, Proserpio V, et al. Accounting for technical noise in single-cell RNA-seq experiments. Nature methods. 2013;10(11):1093.

28. Grün D, Kester L, Van Oudenaarden A. Validation of noise models for single-cell transcriptomics. Nature methods. 2014;11(6):637.

29. Kiselev VY, Kirschner K, Schaub MT, Andrews T, Yiu A, Chandra T, et al. SC3: consensus clustering of single-cell RNA-seq data. Nature methods. 2017;14(5):483–486.

30. Yip SH, Wang P, Kocher JPA, Sham PC, Wang J. Linnorm: improved statistical analysis for single cell RNA-seq expression data. Nucleic acids research. 2017;45(22):e179–e179.

